# An lp17-encoded small non-coding RNA with a potential regulatory role in mammalian host adaptation by the Lyme disease spirochete

**DOI:** 10.1101/2020.11.06.371013

**Authors:** Michael A. Crowley, Troy Bankhead

## Abstract

The bacterial agent of Lyme disease, *Borrelia burgdorferi,* relies on an intricate gene regulatory network to transit between the disparate *Ixodes* tick vector and mammalian host environments. We recently reported that a *B. burgdorferi* mutant lacking an intergenic region of lp17 displayed attenuated murine tissue colonization and pathogenesis due to altered antigen expression. In this study, a more detailed characterization of the putative regulatory factor encoded by the region was pursued through genetic complementation of the mutant with variants of the intergenic sequence. *In cis* complemented strains featuring mutations aimed at eliminating potential BBD07 protein translation were capable of full tissue colonization, suggesting that the region encodes an sRNA. *In trans* complementation resulted in elevated transcription levels and was found to completely abolish infectivity in both immunocompetent and immunodeficient mice. Quantitative analysis of transcription of the putative sRNA by wild type *B. burgdorferi* showed it to be highly induced during murine infection. Lastly, targeted deletion of this region resulted in significant changes to the transcriptome, including genes with potential roles in transmission and host adaptation. The findings reported herein strongly suggest that this lp17 intergenic region encodes for an sRNA with a critical role in the gene regulation required for adaptation and persistence of the pathogen in the mammalian host.

**Author Summary:** Lyme disease continues to emerge as a devastating infection that afflicts hundreds of thousands of people annually in the United States and abroad, highlighting the need for new approaches and targets for intervention. Successful development of these therapies relies heavily on an improved understanding of the biology of the causative agent, *Borrelia burgdorferi.* This is particularly true for the critical points in the life cycle of the pathogen where it must transition between ticks and mammals. Variation in the levels of bacterial gene expression is the lynchpin of this transition and is known to be driven partly by the activity of regulatory molecules known as small non-coding RNAs (sRNAs). In this work, we characterize one of these sRNAs by providing experimental evidence that the transcribed product does not code for a protein, by testing the effects of its overproduction on infectivity, and by interrogating whether its activity causes changes in expression levels of genes at the level of transcription. The findings of this study provide further evidence that regulatory sRNA activity is critical for transmission and optimal infectivity of *B. burgdorferi* and contribute to the recently growing effort to attribute specific roles to these important molecules in the context of Lyme disease.

## Introduction

Lyme borreliosis is an emerging disease with no reliable vaccine that affects hundreds of thousands of people each year in the United States (1). Unmitigated infection with the causative spirochetal bacterium, *Borrelia burgdorferi*, can result in debilitating clinical manifestations in humans including arthritis, carditis, and neurological disorders (2–5). The pathogen is transmitted to humans and other susceptible animals during a bloodmeal taken by feeding *Ixodes* species of ticks.

During transmission from the *Ixodid* tick vector to a mammalian host, *B. burgdorferi* undergoes a drastic shift in gene expression in response to the biophysiochemical disparity between the two environments. This shift in gene expression is controlled in part by the Rrp2-RpoN-RpoS two-component regulatory system, and has been shown to be triggered by changes in temperature, pH, CO2 concentration, host immune pressures, and nutrient availability among other factors (6–12). When expressed, the RpoS alternative sigma factor promotes expression of a subset of genes that facilitate spirochete survival in mammalian hosts (13). RpoS-activated genes include those that encode surface lipoproteins required for host interaction such as complement binding proteins (CRASPs) and extracellular matrix binding proteins (BBK32, P66), proteins essential for surviving innate and adaptive immune responses (OspC and VlsE, respectfully), and metabolic genes required to sustain the bacterium’s auxotrophic nature in the host (PncA) (14–27). Importantly, activation of the RpoS regulon also results in the repression of tick- and transmission-phase associated genes which, if left unrepressed, can result in attenuated ability of spirochetes to establish persistent murine infection (28).

To date, study of the transcriptomic shift resulting from RpoS activation has been primarily focused on two aspects: 1) the bacterial, vector, and host factors that induce gene expression changes, and 2) the end effects of these changes on the transcriptome and proteome of the bacterium under various conditions. The intermediate molecular factors bridging Rrp2-RpoN-RpoS activation and its downstream effects remain largely undetermined. Attempts to bridge this knowledge gap have turned in part towards the investigation of regulatory small non-coding RNA (sRNA) activities in this emerging pathogen.

It was originally thought that the *B. burgdorferi* genome encoded only a few sRNA molecules, and lacked any orthologues of sRNA-associated proteins such as the RNA chaperone protein Hfq (29). However, a series of more recent studies have highlighted the importance of sRNAs in the Lyme disease pathogen as robust facilitators of genetic regulation [Reviewed in (30)]. Lybecker and Samuels demonstrated that *B. burgdorferi* encodes an Hfq orthologue (Hfq_Bb_) that it is required for the function of DsrA, an sRNA important for the translation of RpoS in a temperature-dependent manner (31,32). While other activities exist, this and many other regulatory sRNAs characterized thus far in other pathogenic bacteria operate at the post-transcriptional level through base pairing interactions with target mRNA(s). Several mechanisms have been reported, such as translational activation that includes protection of the transcript from degradation and exposure of the ribosomal binging site following sRNA-mRNA complex formation. Conversely, translational repression can occur, involving destabilization of the transcript or steric prohibition of ribosome binding (33–35).

A recent study by Popitsch *et al* probed the temperature-dependent sRNA transcriptome of *B. burgdorferi* and detected a number of sRNAs that are up-regulated at 37°C compared to room temperature *in vitro,* which is suggestive of a role in mammalian infection (36). Similar work analyzed the stringent response-regulated sRNA transcriptome using a *relB* mutant clone (37). In both studies, an lp17-encoded sRNA was detected and denoted SR0726. This respective RNA transcript mapped to an intergenic space that was previously annotated as a predicted ORF and denoted *bbd07*. An RNA transcript from this same region was also detected in a pair of earlier studies investigating Rrp2-, RpoN-, and RpoS-dependent genes, where its expression was shown to be highly dependent on an intact alternative sigma factor pathway (38, 39).

We recently demonstrated that an lp17 left-end deletion mutant lacking this region displays attenuated murine tissue colonization and pathogenicity, which was ultimately attributed to a 317 bp intergenic region encompassing the *bbd07/* SR0726 locus (40). The tissue colonization defect was not observed during infection of SCID mice, indicating a potential role for this locus in avoidance of adaptive immunity. Further study provided evidence that deletion of this region results in dysregulated antigen expression, implicating the gene product as a regulatory factor.

In the current study, we aimed to test the hypothesis that the *bbd07/*SR0726 region of lp17 encodes an intergenic regulatory sRNA that participates in the transcriptome and proteome shift required for spirochetes to adapt to the mammalian host environment. To do this, BBD07 protein production was first ruled out experimentally through non-native *bbd07/*SR0726 complementation in the previously described lp17 left-end mutant (40). Then, the effects of *bbd07*/SR0726 overexpression were studied in the context of murine infection and antigen expression. These tests revealed that spirochetes expressing high levels of sRNA *bbd07*/SR0726 cannot infect mice, and that this phenotype may be independent of altered *in vitro* antigen expression. Next, the expression of this sRNA was quantified under *in vivo* conditions, which confirmed its putative involvement in the mammalian portion of the enzootic cycle. Finally, a targeted knockout clone of *bbd07*/SR0726 was generated and used for RNA-seq analysis, which revealed that its activity has effects on transcript levels in addition to the previously observed effects on the *in vivo*-expressed antigenic proteome. This work represents a significant step forward towards understanding the critical role that this sRNA has in host adaptation by the Lyme disease pathogen.

## Results

### Mutations that disrupt potential BBD07 protein production do not perturb the functionality of the gene product during murine infection

The recent detection of a putative sRNA encoded within the intergenic space of lp17 containing the discontinued *bbd07* ORF annotation (NC_001849.1 [discontinued]), coupled with numerous failed attempts to detect a BBD07 protein product, provided a strong indication that the *bbd07*/SR0726 locus encodes an sRNA. However, more conclusive results were needed to definitively identify the gene product as an sRNA.

To further rule out the possibility of BBD07 protein production, two *in cis* complement strains were generated in an lp17 mutant strain lacking a region containing *bbd01-bbd07* that is incapable of murine heart tissue colonization (Δ1-7; (40)). These two genetically complemented strains harbored a *bbd07*/SR0726 copy containing either a premature stop codon or a disrupted start codon, and were denoted Comp7Nstop_*c*_ and Comp7NΔstart_*c*_, respectively (Table 1). The rationale was that if the region encodes a protein, then these strains would not be capable of translating the functional native protein, leading to a defective tissue colonization phenotype comparable to the Δ1-7 mutant strain. Alternatively, if this region encodes an sRNA, then these mutations would be expected to have no effect on functionality, and thus the strains would exhibit tissue colonization levels comparable to the wild type control and native *cis* complement (Comp7N_*c*_) strains.

**Table 1.**
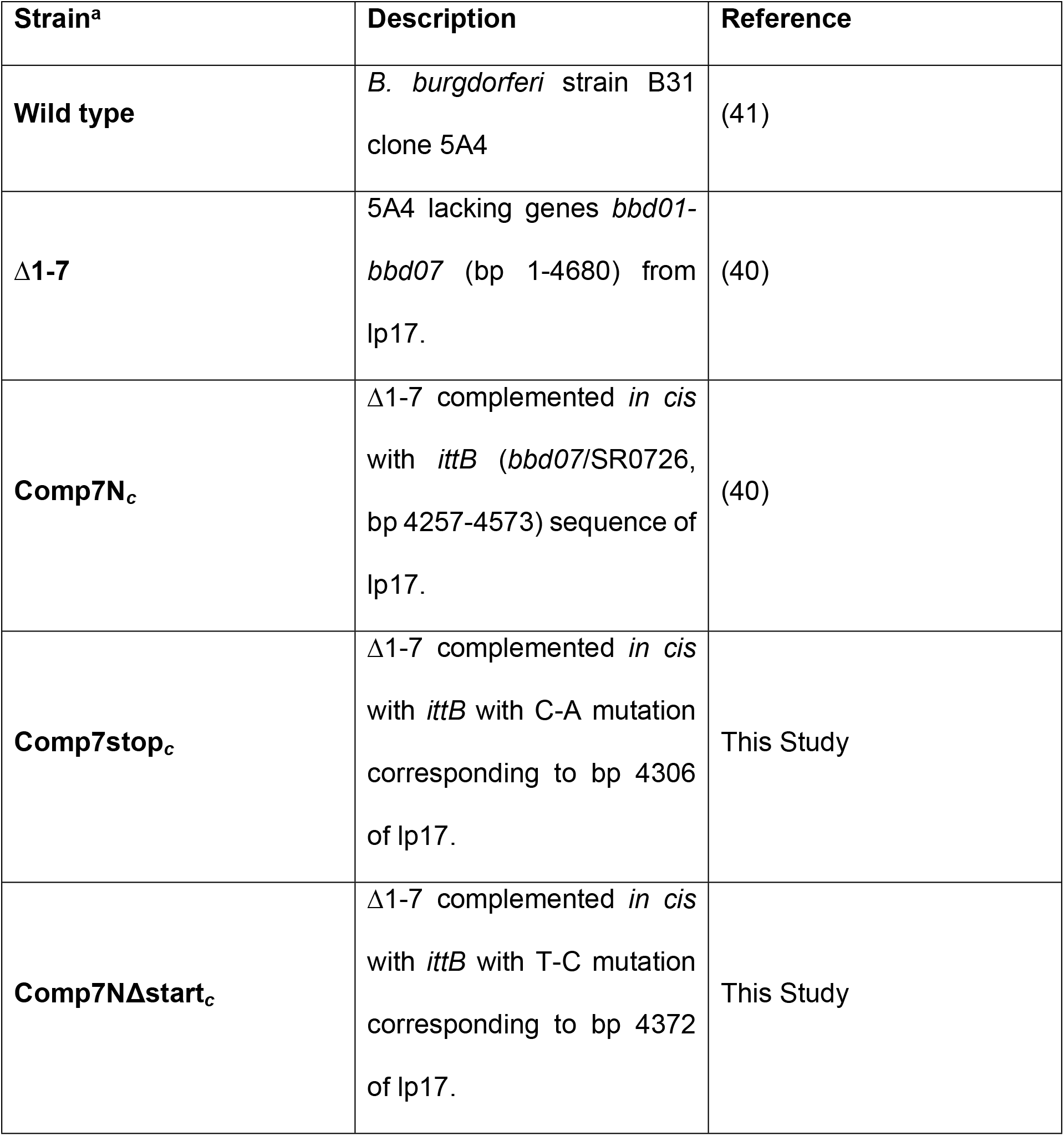
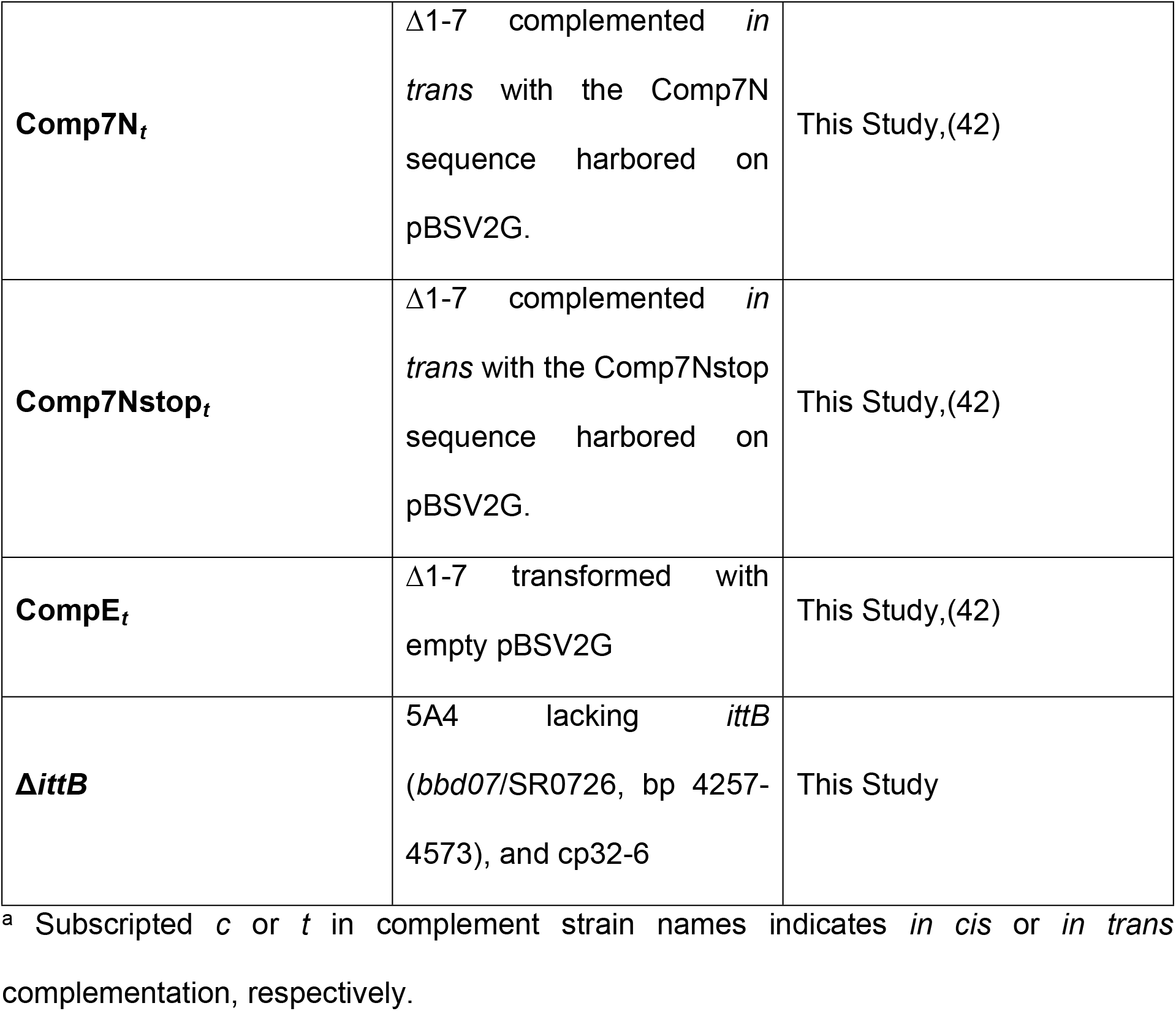
*B. burgdorferi* strains used in this study.

| Strain<sup>a</sup> | Description | Reference |
| --- | --- | --- |
| **Wild type** | *B. burgdorferi* strain B31 clone 5A4 | (41) |
| **Δ1-7** | 5A4 lacking genes *bbd01-bbd07* (bp 1-4680) from lp17. | (40) |
| **Comp7N<sub>c</sub>** | Δ1-7 complemented *in cis* with *ittB* (*bbd07*/SR0726, bp 4257-4573) sequence of lp17. | (40) |
| **Comp7stop<sub>c</sub>** | Δ1-7 complemented *in cis* with *ittB* with C-A mutation corresponding to bp 4306 of lp17. | This Study |
| **Comp7NΔstart<sub>c</sub>** | Δ1-7 complemented *in cis* with *ittB* with T-C mutation corresponding to bp 4372 of lp17. | This Study |
| **Comp7N<sub>t</sub>** | Δ1-7 complemented *in trans* with the Comp7N sequence harbored on pBSV2G. | This Study,(42) |
| **Comp7Nstop<sub>t</sub>** | Δ1-7 complemented *in trans* with the Comp7Nstop sequence harbored on pBSV2G. | This Study,(42) |
| **CompE<sub>t</sub>** | Δ1-7 transformed with empty pBSV2G | This Study,(42) |
| **ΔittB** | 5A4 lacking *ittB* (*bbd07*/SR0726, bp 4257-4573), and cp32-6 | This Study |

<sup>a</sup> Subscripted *c* or *t* in complement strain names indicates *in cis* or *in trans* complementation, respectively.

Single nucleotide mutations were made in frame as defined by the previously predicted TTG start codon at bp 4373 of *B. burgdorferi* B31A lp17(NC_001849). For Comp7Nstop*c*, the stop codon was generated via a cytosine to adenine mutation corresponding to bp 4306 of the annotated lp17 DNA sequence. For Comp7NΔstart_*c*_, a thymine to cytosine mutation was introduced in the middle nucleotide of the previously predicted start codon that corresponded to bp 4372 of the annotated lp17 DNA sequence (Fig. 1A). Neither mutation was predicted to significantly perturb the putative RNA secondary structure that could be important for the potential function of the *bbd07*/SR0726 sRNA. To generate the clones, TOPO-derived plasmids harboring the 317 bp sequences were used to transform Δ1-7 *B. burgdorferi* cells, and transformants were screened for insertion of the altered sequences in lp17. PCR verification of these clones demonstrated successful *bbd07*/SR0726 complementation as shown in Figure 1B. Sequencing was performed to verify the single base pair mutations in each construct, and isogeneity to the parent clone was confirmed by multiplex PCR for native plasmid content (data not shown) (43).

**Fig 1.**
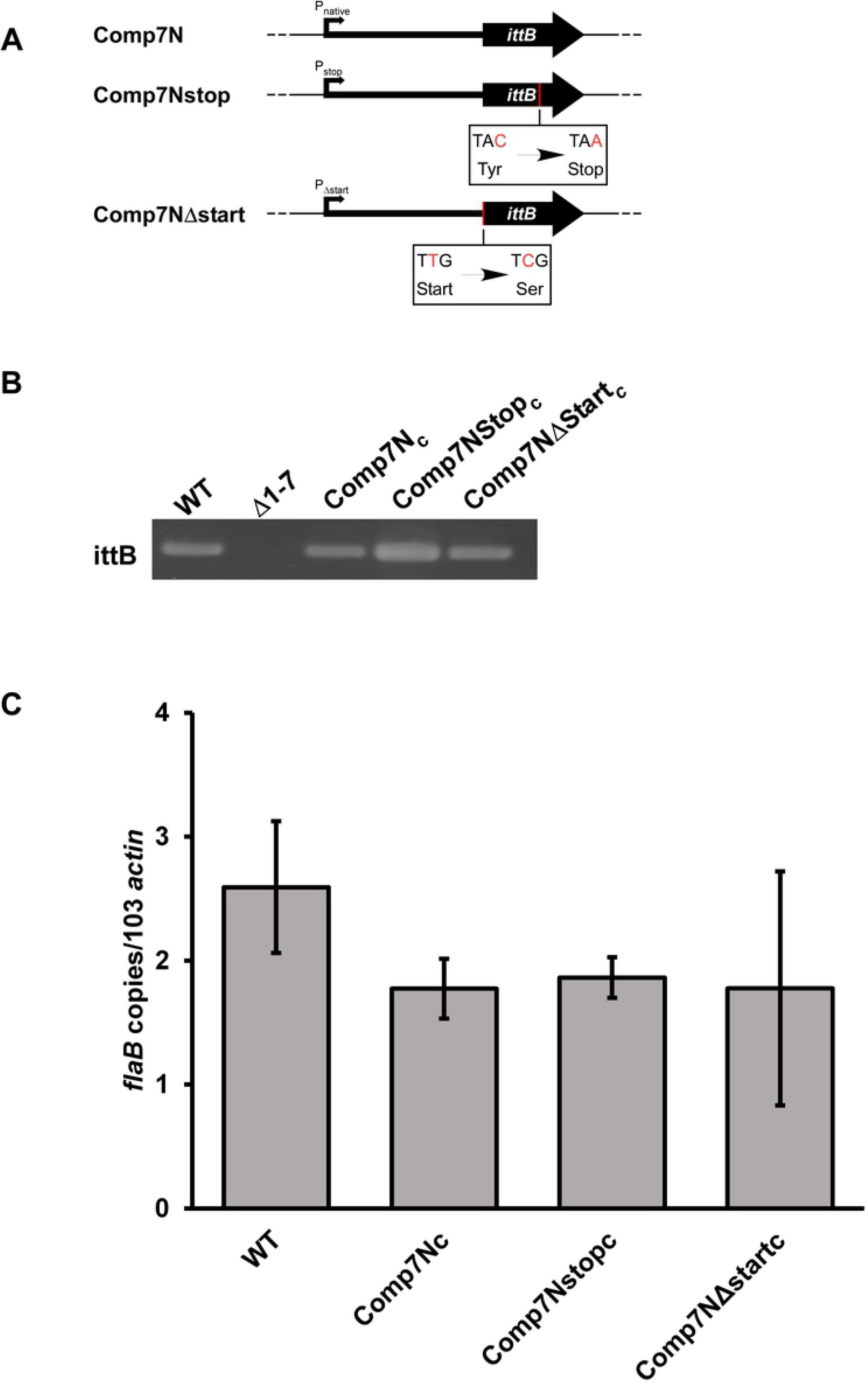
Neither introduction of a stop codon nor disruption of potential start codon abrogates the function of the *bbd07*/SR0726 gene product. A) Diagrams indicating where mutations were made with respect to the discontinued ORF annotation. Mutated base pairs are indicated with red text. B) PCR verifications of successful complementation using P1050/P1051. C) Mice were inoculated subcutaneously with indicated strains at a dose of 5×10^3^ total spirochetes. qPCR analysis was used to assess bacterial burden in heart tissues harvested from mice infected with the indicated strains for 28 days. Reactions were performed in duplex with probes/primers for *flaB* (P199, P200, P201) and mouse *actin* (P202, P203, P1086). Data bars indicate the average number of *flaB* copies per 10^3^ *actin* copies in heart tissues from 5 mice. Students t-test was performed to assess significance of differences in heart tissue colonization between each recombinant strain and the wild type, p>0.05. Error bars indicate SEM for each group (n=5).

To determine if the altered *bbd07*/SR726 sequences were able to rescue the mutant phenotype, the capacity for heart tissue colonization of immunocompetent mice was selected for use as a readout. We reasoned that this was a reliable indicator of gene functionality due to the fact that the Δ1-7 mutant has been previously shown to be unable to colonize heart tissue in immunocompetent mice (40). Groups of five C3H mice each were needle inoculated (5×10^3^ total spirochetes) with either wild type or one of the various complement clones. Successful establishment of infection was monitored weekly by culturing blood and ear tissues to visually verify the presence of spirochetes by dark field microscopy (Table 2). All mice in the experiment displayed spirochetemia at 7 days post infection and remained infected for the entire 28-day duration of the experiment. All ear tissue samples on day 14 post infection from Comp7Nstop_*c*_- and Comp7NΔstart_*c*_-infected mice yielded spirochete growth in culture. At day 28 post infection, heart tissue was collected from each mouse and divided in half longitudinally. One half of each heart was deposited into culture media, while the other half was snap-frozen for DNA extraction and qPCR to determine bacterial burden.

**Table 2.** Tissue colonization capacity of *bbd07* complement strains.

| Strain | Time point/tissue |  |  |  |
| --- | --- | --- | --- | --- |
|  | Day 7/ blood | Day14/ ear | Day 21/ ear | Day 28/ heart |
| **Wild type** | 5/5 | 5/5 | 5/5 | 5/5 |
| **$\Delta 1-7$** | 13/13 | ND | ND | 0/13* |
| **Comp7N<sub>c</sub>** | 5/5 | 5/5 | 5/5 | 5/5 |
| **Comp7Nstop<sub>c</sub>** | 5/5 | 5/5 | 5/5 | 5/5 |
| **Comp7NDstart<sub>c</sub>** | 5/5 | 5/5 | 5/5 | 5/5 |

\* Significant difference compared to wild type and each complemented strain as determined by Fisher's exact test ( $p < 0.05$ ). ND; not determined.

Comp7Nstop_c_ and Comp7NΔstart_c_ spirochetes were recovered from heart tissue to the same extent as the wild type and Comp7N_c_ strains (Table 2). Moreover, qPCR analysis of total DNA from the tissues showed no statistically significant differences in spirochete burden between any of the tested strains (Fig. 1C). These combined findings demonstrate that neither introduction of a premature stop codon nor the destruction of the potential start codon affect the functionality of the gene product in the context of heart tissue colonization, further suggesting that the *bbd07*/SR0726 region encodes a sRNA and not a protein product. As this sRNA is encoded on lp17 and has a phenotype similar to *ittA* (infection-associated and tissue-tropic sRNA A) recently reported by another group (44), we have chosen to designate the functional RNA product of the *bbd07*/SR0726 locus as *ittB* (infection-associated and tissue-tropic sRNA B).

### Overexpression of *ittB* is deleterious to murine host infection

Our laboratory recently reported that absence of the region encoding *ittB* affects antigen expression *in vivo,* which could potentially explain the observed attenuation in tissue colonization and pathogenesis (40). Thus, it was hypothesized that *ittB* overexpression may lead to altered murine infectivity due to abnormal antigen production. *B. burgdorferi* has been shown to maintain the pBSV2G shuttle vector at a higher copy number than its native plasmids, and expression levels of genes complemented on this vector can be elevated compared to wild type (42). Thus, *in trans* complemented strains that harbored *ittB* gene copies on pBSV2G were generated in the Δ1-7 mutant background to assess the effects of *ittB* overexpression. Complementation was performed by transforming Δ1-7 cells with pBSV2G carrying either a wild-type copy of *ittB* (Comp7N_*t*_) or a copy containing an early stop codon (Comp7Nstop_*t*_). An empty vector control strain harboring pBSV2G without an *ittB* gene copy was also generated (CompE_*t*_). PCR verification of *ittB* presence or absence in these strains is illustrated in Figure 2A, and isogeneity to the parent strain was confirmed via multiplex PCR (data not shown).

**Fig 2.**
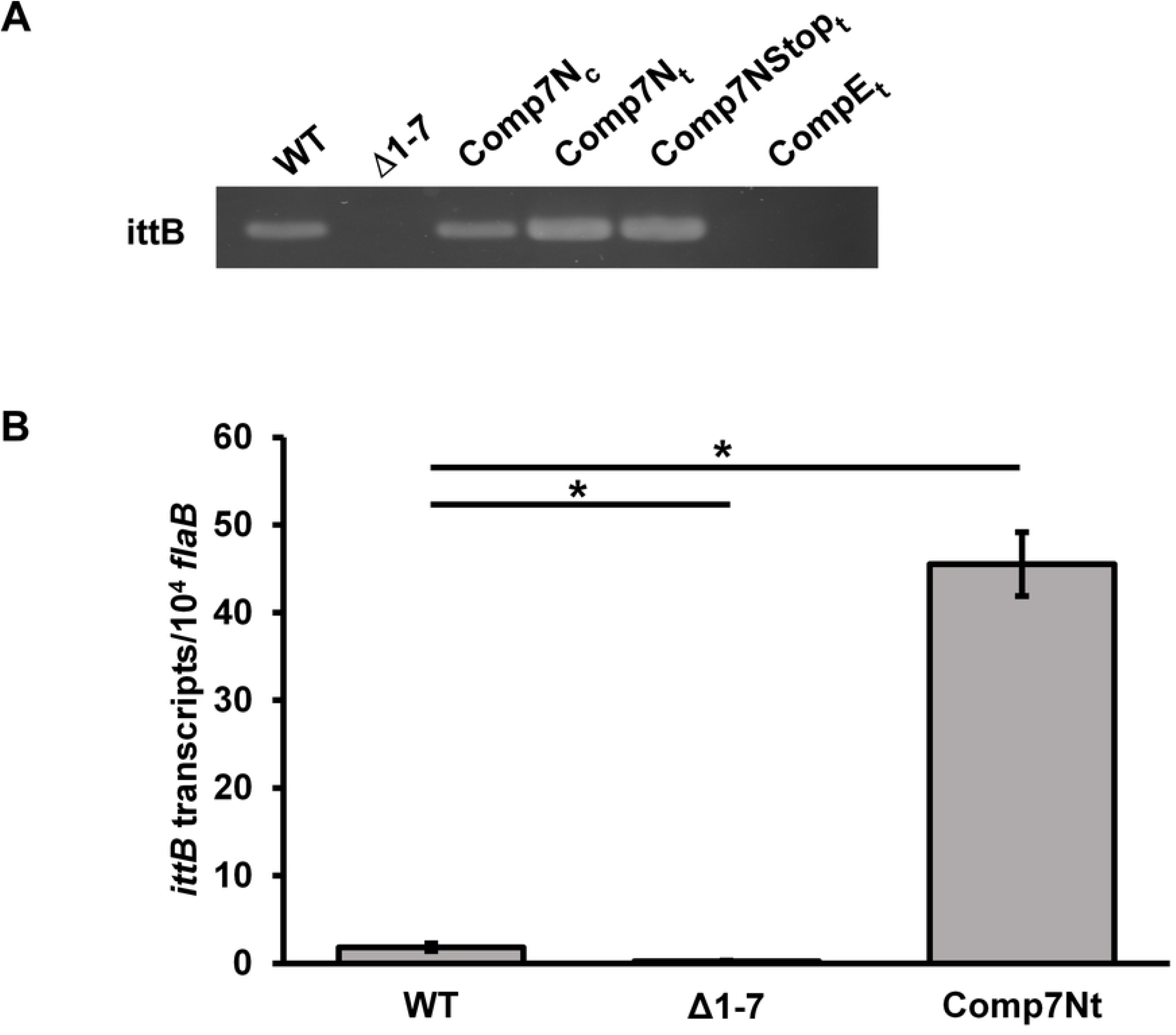
*In trans* complementation of *ittB* results in elevated transcription. A) PCR verifications of successful complementation using P931/P932. B) Assessment of *ittB* transcript levels by wild-type, Comp7N_*c*_, and Comp7N_*t*_ in vitro via qRT-PCR. Triplicate cultures were grown to late log-phase (10^8^ spirochetes mL^-1^). Reactions were performed in duplex with primers for *ittB* (P1001/P1003) and *flab* (P411/P412). Data bars indicate the average number of *ittB* copies per 10^4^ *flaB* copies, with lines indicating standard error mean for each group of replicates. Data from negative control reactions using Δ1-7 cDNA is included to illustrate background signal in the assay. Students t-test was performed to assess significance of the differences in *ittB* transcription between each strain, *p≤0.05.

To determine the relative *in vitro* expression levels of *ittB* in the wild type and Comp7N_*t*_, qRT-PCR analysis was performed using cDNA derived from triplicate cultures of wild type, Δ1-7, and Comp7N_*t*_ grown to late log-phase (1×10^8^ spirochetes ml^-1^). As shown in Figure 2B, a ~24-fold increase in *ittB* expression by Comp7N_t_ spirochetes compared to the wild type was observed. Wild-type expression of *ittB* was found to be low, producing less than 2 *ittB* copies per 10^4^ *flaB* copies. These results indicate that *ittB* may be tightly regulated, and that residence of the *ittB* sequence on a high-copy vector may disrupt this regulation that results in elevated *in vitro* transcription.

To assess the effects of high copy *ittB* expression on murine infection, groups of five C3H mice each were needle inoculated (5×10^3^ total spirochetes) with wild type, Comp7N_*t*_, Comp7Nstop_*t*_, or CompE_*t*_ spirochetes. Infection was monitored weekly by culturing of tissues as before. Interestingly, while infection was established in all mice inoculated with the wild type or CompE_t_ control clone, neither Comp7N_*t*_ nor Comp7Nstop_*t*_ spirochetes could infect any mice (Table 3). To ensure reproducibility, a second independent infection assay with the Comp7N_*t*_ and Comp7Nstop_*t*_ clones was conducted with the same results. Next, groups of three C3H mice each were inoculated with 10^4^, 10^5^, or 10^6^ Comp7N_*t*_ spirochetes alongside three mice infected with 10^4^ wild type spirochetes in order to test if the infectivity defect associated with *ittB* overexpression was dependant on the inoculum dosage. While all wild type-inoculated mice became infected, none of the mice inoculated with Comp7N_*t*_ became infected despite a higher inoculum dosage, indicating that this phenotype is not dose dependent. To determine whether the lack of infectivity exhibited by these strains in C3H mice was due to an inability to evade the adaptive immune response, Comp7N_t_ and Comp7Nstop_t_ spirochetes were used to inoculate groups of five immunodeficient SCID mice. Neither strain was able to establish infection in any of the mice tested during the 28-day experiment (Table 4). Together, these findings suggest that high-copy expression of *ittB* renders spirochetes unable to establish murine infection, even in the absence of an adaptive immune response.

**Table 3.**
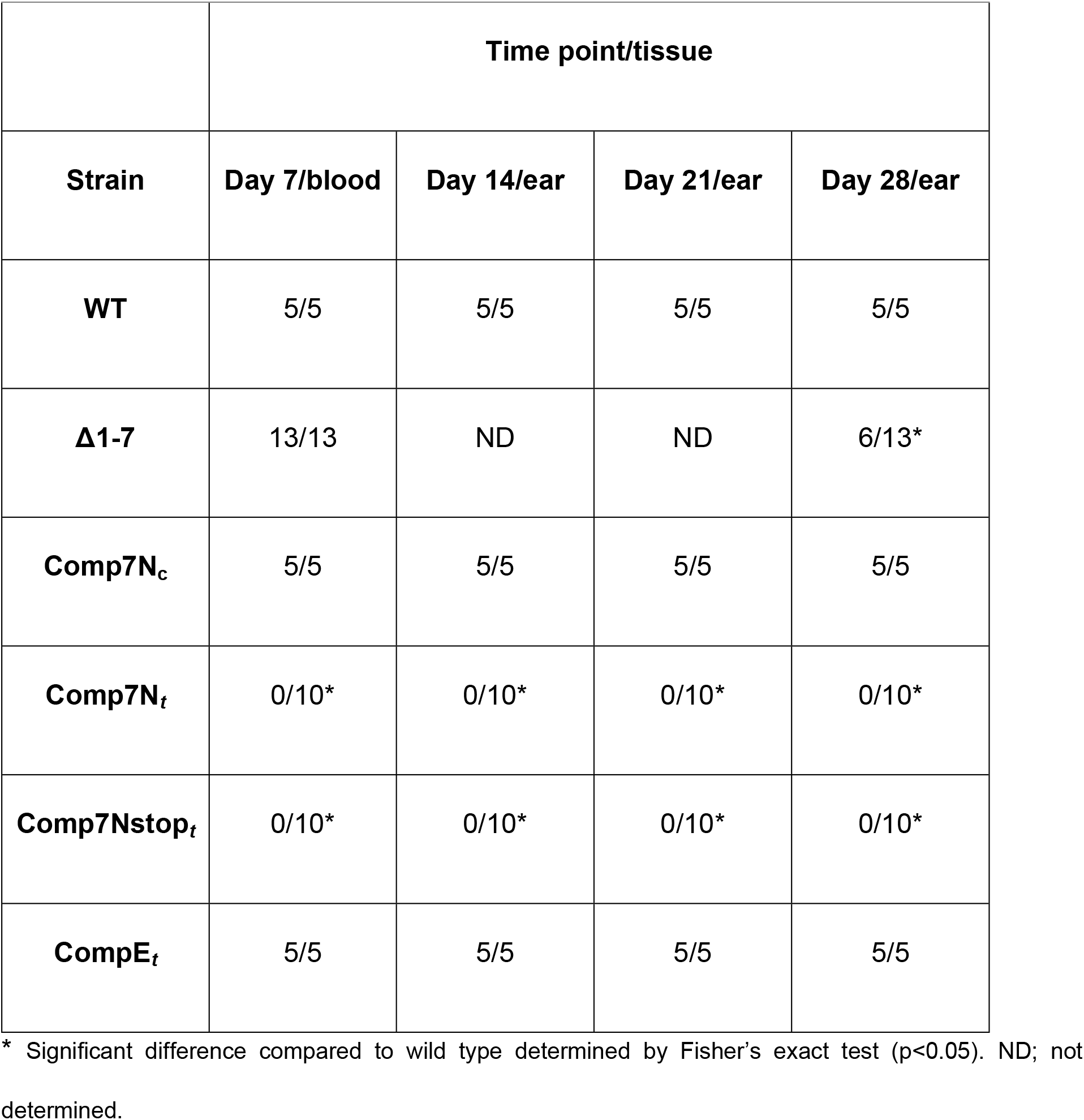
Infectivity of *in trans* complement strains in C3H mice.

| Strain | Time point/tissue |  |  |  |
| --- | --- | --- | --- | --- |
|  | Day 7/blood | Day 14/ear | Day 21/ear | Day 28/ear |
| **WT** | 5/5 | 5/5 | 5/5 | 5/5 |
| **$\Delta 1-7$** | 13/13 | ND | ND | 6/13* |
| **Comp7N<sub>c</sub>** | 5/5 | 5/5 | 5/5 | 5/5 |
| **Comp7N<sub>t</sub>** | 0/10* | 0/10* | 0/10* | 0/10* |
| **Comp7Nstop<sub>t</sub>** | 0/10* | 0/10* | 0/10* | 0/10* |
| **CompE<sub>t</sub>** | 5/5 | 5/5 | 5/5 | 5/5 |

\* Significant difference compared to wild type determined by Fisher's exact test ( $p < 0.05$ ). ND; not determined.

### High-copy expression of *ittB* does not alter antigen expression *in vitro*

Recently, we showed that the murine tissue colonization defect and attenuated pathogenesis exhibited by the Δ1-7 mutant during infection was associated with dysregulated antigen expression (40). Considering this, along with the findings that the *in trans ittB* complement strains were non-infectious in mice, it seemed possible that overexpression of *ittB* might also result in dysregulated antigen expression that in turn renders those strains non-infectious. Because *ittB* transcription by Comp7N_*t*_ spirochetes is highly elevated during *in vitro* cultivation (Fig. 2B), we predicted that any potential resultant protein expression changes would be observable under the same conditions.

To assess the effects of *ittB* overexpression on overall antigen production, cultures of wild type, Δ1-7, Comp7N_*c*_, Comp7N_*t*_, and CompE_*t*_ spirochetes were grown in triplicate under the same conditions used to quantify *ittB* transcript levels in the Comp7N_*t*_ strain. Protein lysates (10^9^ total cells) of each strain were subjected to Western blot analysis using murine immune sera harvested from mice that had been infected with wild type *B. burgdorferi* for 28 days. Surprisingly, no notable differences could be observed between the *in vitro* antigenic profiles of the strains tested (Fig. 3). This result is suggestive that the non-infectious phenotype exhibited by Comp7N_*t*_ may not be due to dysregulated antigen expression.

**Fig 3.**
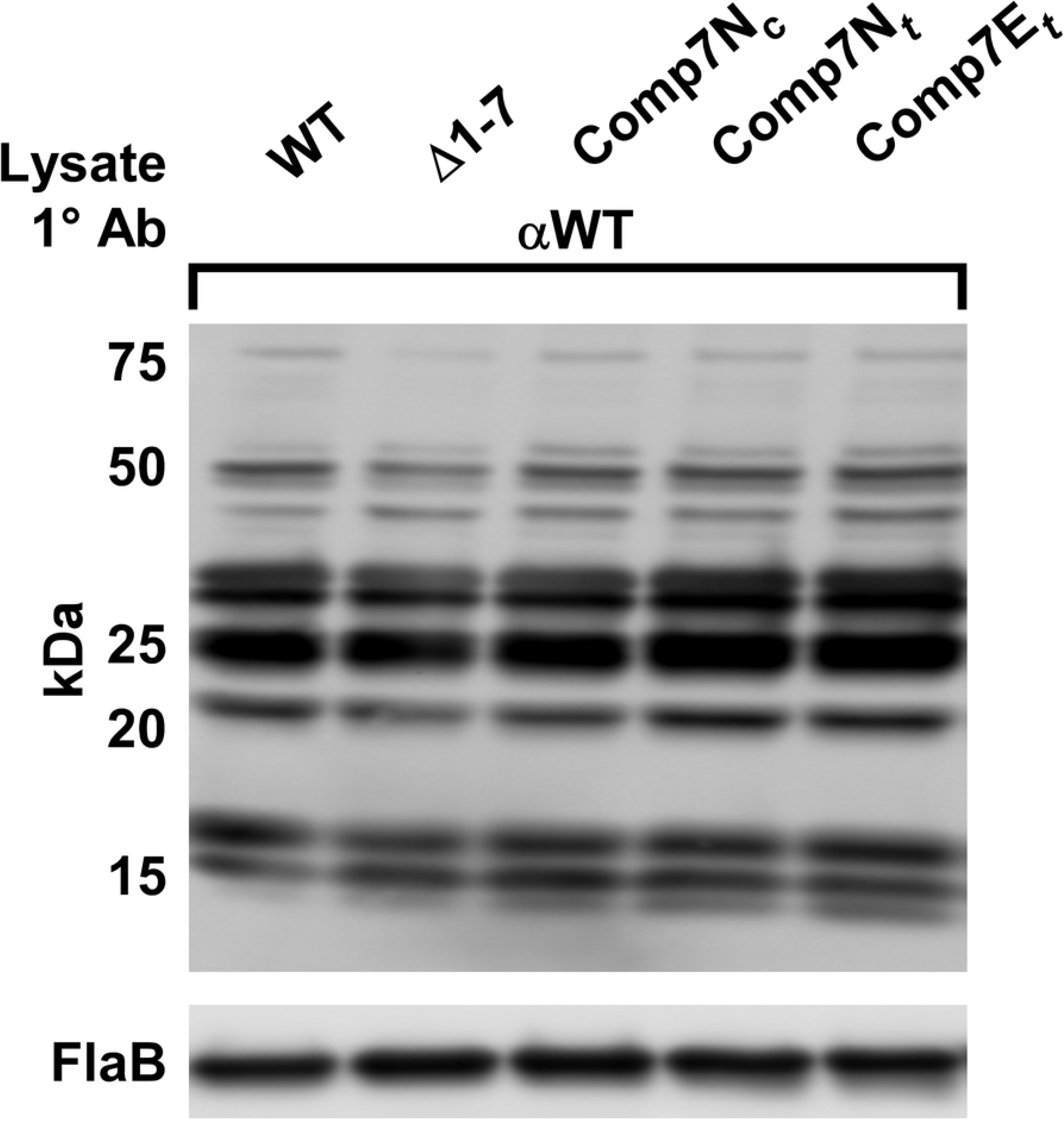
High-copy expression of *ittB* does not result in dysregulation of antigen expression *in vitro.* Wild type, Δ1-7, Comp7N_c_, Comp7N_t_, and Comp7E_t_ strains were grown *in vitro* in triplicate under the same conditions used for *in vitro ittB* transcript quantification. Bacterial lysates (10^9^ cells per lane) were Western blotted with serum collected from C3H mice infected with wild type for 28 days. FlaB protein from these lysates was probed with polyclonal anti-FlaB antibodies as a loading control. Approximate molecular weights are indicated to the left.

### Expression of *ittB* is induced during murine infection

Transcription of the *ittB* locus was previously shown to increase during *in vitro* growth at 37°C compared to 23°C, suggesting that it is upregulated during transmission and/or infection of the mammalian host (36). To verify that *ittB* transcription is induced in the mammalian host, and to assess any potential expression differences in spirochetes colonizing various tissue sites, *ittB* RNA transcripts were quantified from infected murine tissues by qRT-PCR. Groups of three mice each were infected with 1×10^5^ total wild-type spirochetes, and heart, bladder, and joint tissues were harvested at 28 days post infection for total RNA extraction and subsequent analysis. Relative abundance of *ittB* expression (*ittB* copies per *flab* copies) was calculated for each tissue. Expression of *ittB* was detected in each tissue examined, with an average expression level of approximately 20, 65, and 95 copies of *ittB* per 10^3^ copies of *flab* in spirochetes colonizing either the heart, bladder, or joints, respectively (Fig. 4). These values range from a 100- to 400-fold increase in *ittB* transcription levels relative to that observed for *in vitro* grown wild-type spirochetes (~2 copies of *ittB* per 10^4^ copies of *flab,* see Fig. 2B), supporting the hypothesis that *ittB* transcription is upregulated during mammalian infection. Despite *ittB* transcription being ~100-fold higher in heart-resident spirochetes than those grown *in vitro, ittB* expression levels in these spirochetes were significantly lower than those colonizing joint tissue (p≤0.05). Differences in *ittB* transcription either between heart and bladder or bladder and joint did not reach significance.

**Fig 4.**
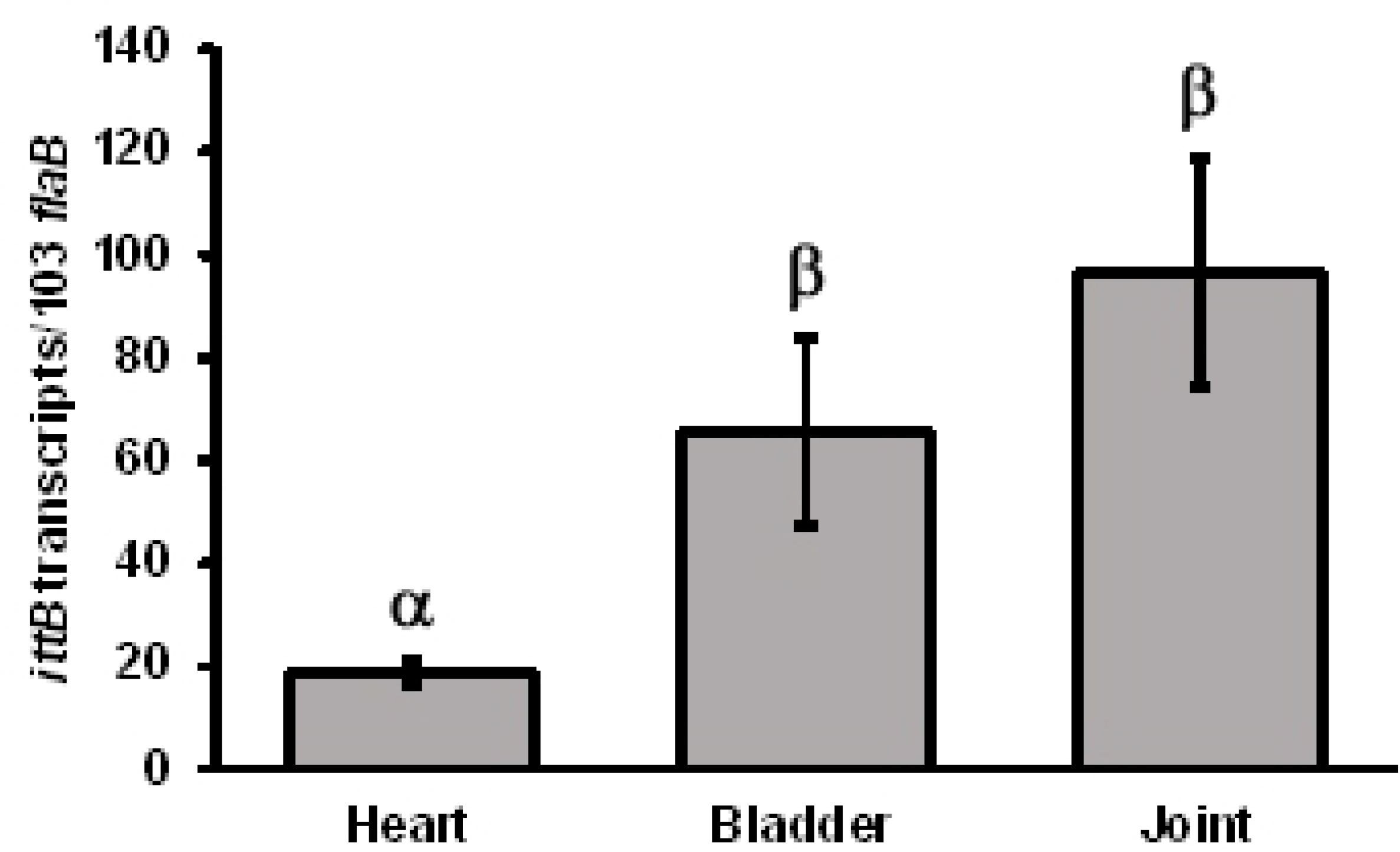
*ittB* expression is induced towards mammalian infection. Three mice were inoculated subcutaneously with 10^5^ wild type spirochetes. Infection was verified as before. Tissues were harvested for RNA extraction on day 28 post-infection. cDNA derived from indicated tissues was used in qRT PCR reactions similar to those used for assessment of *ittB* transcript levels *in vitro*. Error bars indicate SEM among biological replicates (n=3). Symbols α and *β* indicate statistically different groups (*p* < .05) as determined by one-way ANOVA followed by all pairwise multiple comparison (Holm-Sidak).

### Targeted deletion of *ittB* results in an altered transcriptome

To further examine a potential regulatory role for *ittB*, we sought to determine if deletion of *ittB* affects the transcriptome of *B. burgdorferi.* To eliminate any possible secondary effects brought about by the additional loss of *bbd01-bbd06* in the Δ1 −7 mutant clone, it was necessary to generate a knockout clone that only lacked the *ittB* locus. Deletion of the 317 bp region of lp17 encoding *ittB* was performed through allelic exchange using a TOPO-derived plasmid construct containing a gentamycin resistance cassette flanked by upstream and downstream regions of DNA homologous to lp17 (Fig. 5A). A positive clone (Δ*ittB*) was selected and recovered via PCR screening as shown in Figure 5B. The endogenous plasmid profile of this clone was identical to the wild type 5A4 parent strain with the exception of cp32-6, which is not required for infectivity (45). Deletion was further verified by RT-PCR to ensure the absence of *ittB* transcription (Fig. 5C). A complement clone was also generated by transforming *ΔittB* cells with gDNA from a clone harboring an lp17 plasmid in which the *bbd01-bbd03* region was replaced with a kanamycin resistance gene (Δ*ittB*Comp_*c*_, Fig. 5B). *bbd01-bbd03* were previously shown to be dispensable in terms of tissue colonization during murine infection (40). This complementation approach was pursued after numerous attempts to complement the *ittB* region via allelic exchange had failed. Genetic complementation in *Borrelia* strains is well known to be difficult at times, and similar approaches involving plasmid replacement have been utilized in other published studies where traditional complementation proved unsuccessful (46, 47).

**Fig 5.**
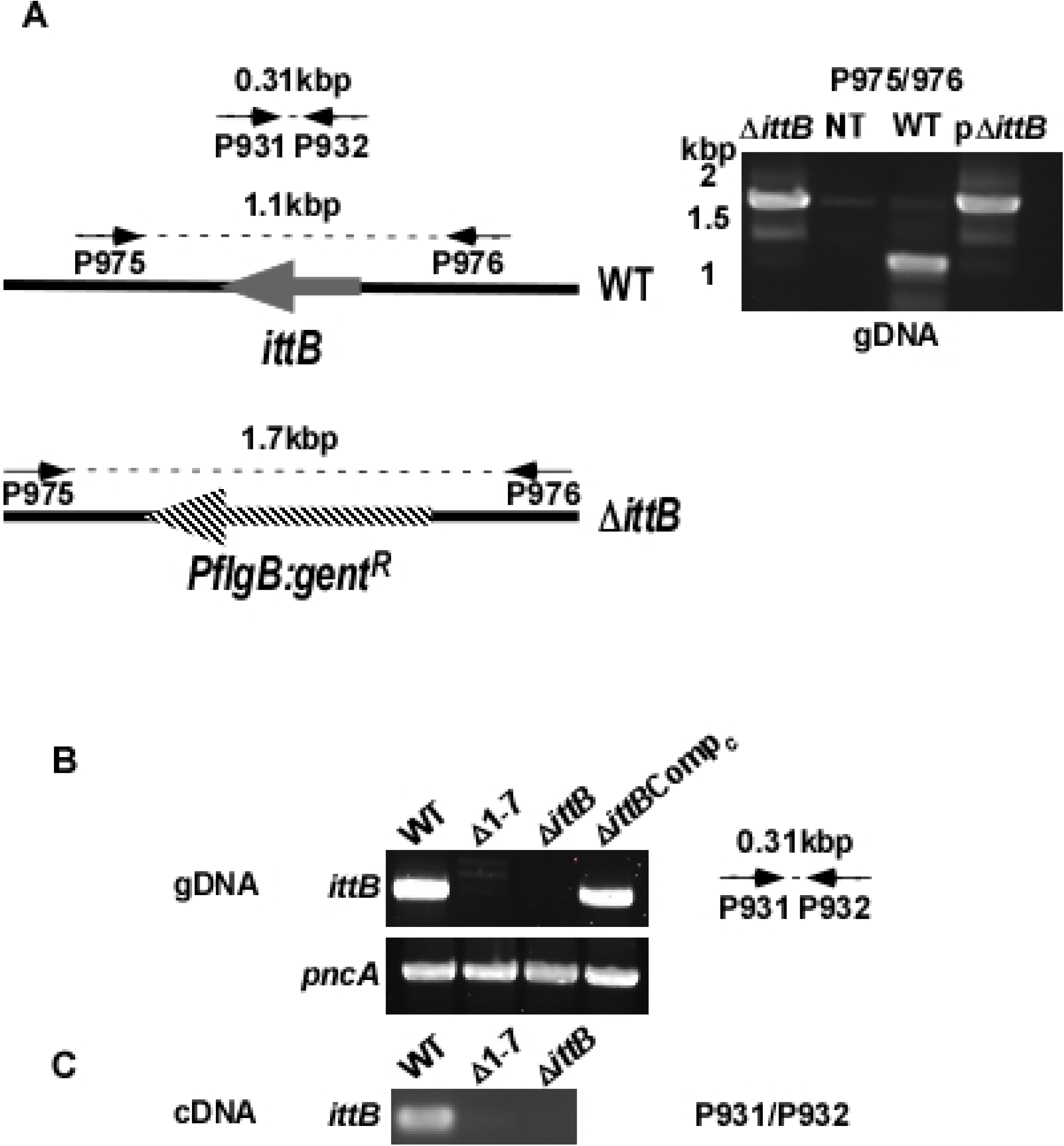
Targeted deletion of *ittB.* A) Schematic illustrating the relevant segment of lp17 in the wild type (WT) and the *ittB* knockout (Δ*ittB*). Screening primer annealing sites and expected amplicon lengths are shown above each schematic with arrows connected by dashed lines. B) PCR confirmations of presence or absence of *ittB* genetic region using P931/P932 and genomic DNA templates from the indicated strains. C) Verification of presence or absence of *ittB* transcription in indicated strains by RT-PCR using P931/P932.

A mouse infection assay was carried out to ensure the newly generated Δ*ittB* mutant strain would exhibit an *in vivo* phenotype similar the Δ1-7 mutant clone (40). Groups of five C3H mice each were needle inoculated with the wild type or Δ*ittB* mutant at a dose of 5×10^3^ total spirochetes. All mice became infected as determined by culture positivity of blood and ear tissue collected weekly following infection. At day 28 post infection, heart, bladder, joint, and ear tissue were harvested and cultured for detection of spirochete growth by dark field microscopy as described before. As was previously observed for Δ1-7, none of the heart tissue cultures from Δ*ittB*-infected mice were positive for spirochete growth, further demonstrating that *ittB* is required for heart tissue colonization in mice (Fig. 6). Unexpectedly, joint tissue colonization was attenuated for Δ*ittB* compared to wild type, with spirochetes being recovered from that tissue in only 2/5 Δ*ittB*-infected mice. Although this difference did not reach statistical significance (p=0.167), it contrasts with previous results observed with the Δ1-7 clone, which was significantly attenuated at day 14 but recovered from joints of all mice at the 28-day time point (40). However, Δ*ittB*Comp_*c*_ spirochetes were found to be able to colonize all tissues in all mice, supporting the validity of the observed phenotype of the targeted *ittB* deletion mutant.

**Fig 6.**
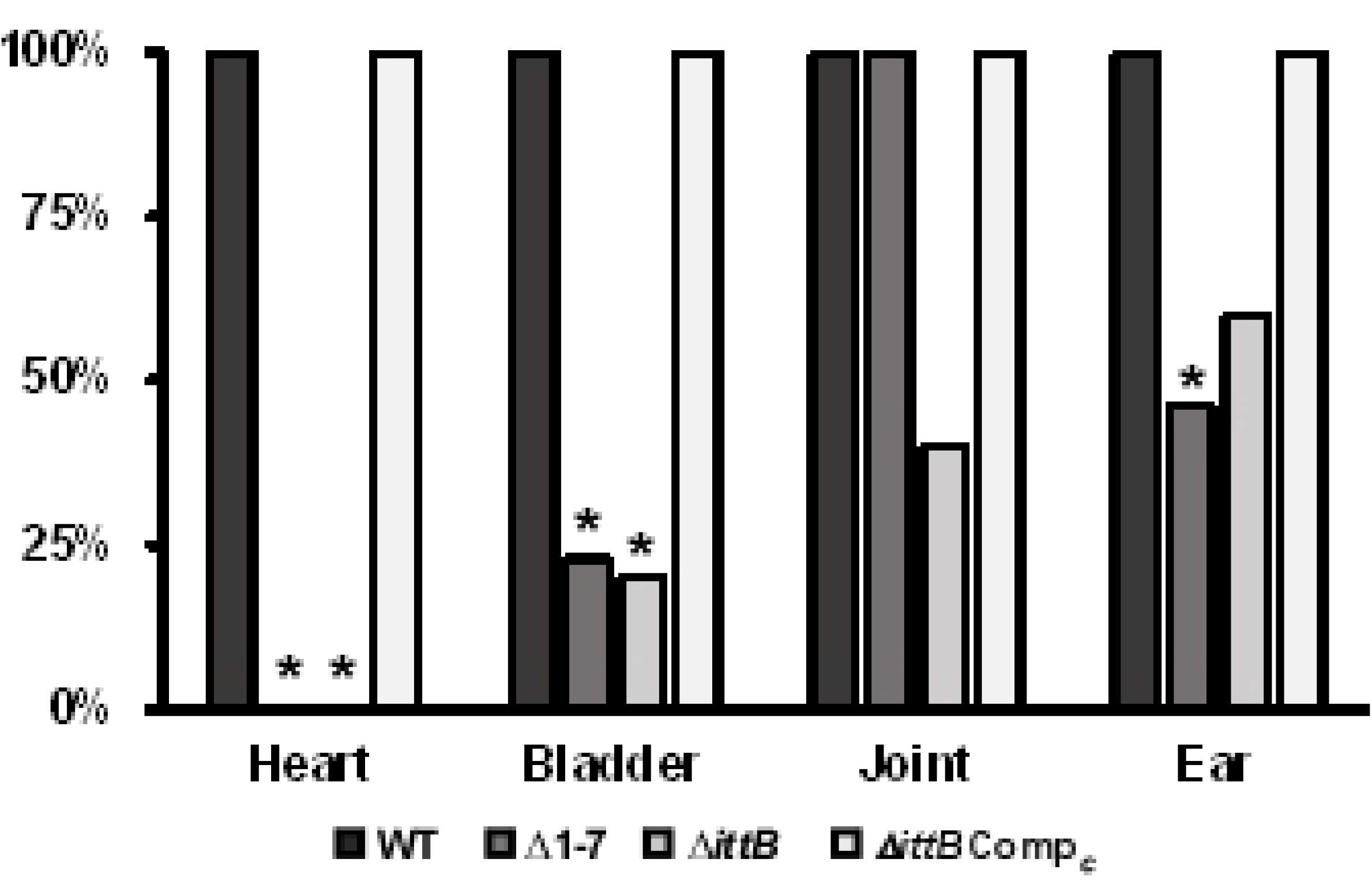
Infectious phenotype of *ΔittB.* Groups of 10, 5, and 4 mice were infected with 5×10^3^ spirochetes of the wild type, *ΔittB,* or Δ*ittB*Comp strains, respectively. Indicated tissues were harvested at day 28 post-infection and cultured in BSK for detection of spirochete growth as described above. Data bars indicate the percentage of mice in each group that were culture positive for spirochete growth in the indicated tissues. Δ1-7 data was transposed from (40). Asterisks indicate statistically significant difference compared with wild type as determined by Fisher’s exact test (p < .05).

As a final confirmation that the *ΔittB* mutant exhibits similar defects to those of the Δ1-7 strain, the *in vivo* antigenic profile of Δ*ittB* was compared to those of the wild type and Δ1-7 strains. This was done by subjecting *in vitro*-grown wild type spirochetes to Western blot analysis using pooled antisera from groups of 3 mice infected for 28 days with either wild type, Δ1-7, or Δ*ittB* spirochetes. Sera from *ΔittB-* and Δ1-7-infected mice displayed detectable reactivity to at least two additional antigens compared to sera from wild type-infected mice (Fig. 7), indicating altered expression of these antigens during murine infection by strains lacking *ittB.*

**Fig 7.**
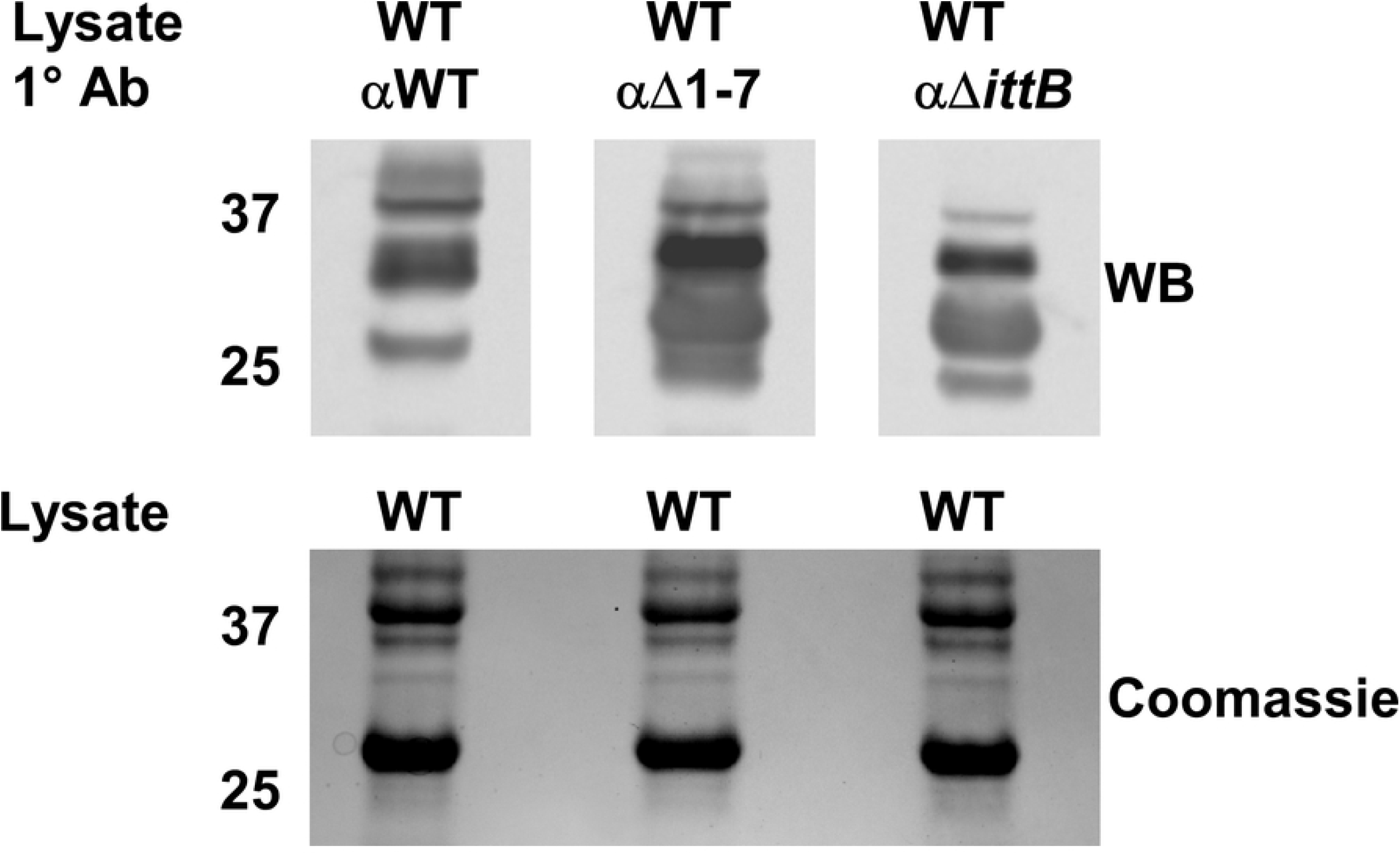
*ΔittB* exhibits erroneous antigen expression *in vivo* comparable to Δ1-7. Antisera were extracted from mice (n=5) infected with the indicated strains for 28 days. The sera were used as the primary antibody in a western blot probing protein lysate of *in vitro* cultivated wild type spirochetes. A gel loaded identically to that used for the Western blot was Coomassie stained to serve as a protein loading control and is shown above the blot.

Finally, the Δ*ittB* mutant was analyzed by RNA-seq to determine whether *ittB* deletion affects the *B. burgdorferi* transcriptome. To accomplish this, triplicate cultures of wild-type or Δ*ittB* spirochetes were grown to a density of 1×10^8^ spirochetes mL^-1^ at 37°C following a temperature shift involving a 1:100 subculture from room temperature cultures. This temperature shift was performed in an attempt to induce *ittB* expression by the wild type (36). RNA from the three pooled cultures was DNase treated and sent to the Novogene Corporation for library preparation, sequencing, and data analysis. Table 4 shows a list of 17 transcripts that were differentially expressed between wild type and Δ*ittB* transcriptomes with a log2-fold change >±2. In total, 92 transcripts were differentially detected with a log_2_-fold change >±1, omitting those transcripts encoded by cp32-6, which is absent in the Δ*ittB* clone. The most significant transcriptional difference observed was in *bb0360,* which was detected approximately 34-fold more from Δ*ittB* compared to wild type (P=3.51×10^-12^). Sultan *et al* demonstrated that this gene is the first in a co-transcribed operon including the subsequent 4 genes *(bb0361-bb0364)* (48). Despite this, none of the other transcripts in the putative polycistron were detected differentially between the strains. *bb0365,* encoded immediately downstream of this operon, was detected approximately 6-fold less from Δ*ittB* compared to wild type. *bb0733* was detected 5.5-fold less from Δ*ittB* compared to wild type and is a well characterized c-di-GMP binding protein. Combined, this data demonstrates that deletion of *ittB* has effects on the *in vitro* transcriptome of *B. burgdorferi,* and that the overall effects of its absence may be diverse.

**Table 4.** Infectivity of *in trans* complement strains in SCID mice.

|  | Time point/tissue |  |  |  |
| --- | --- | --- | --- | --- |
| Strain | Day 7/blood | Day 14/ear | Day 21/ear | Day 28/ear |
| WT | 5/5 | 5/5 | 5/5 | 5/5 |
| $\Delta 1-7$ | 5/5 | 5/5 | 5/5 | 5/5 |
| Comp7N<sub>t</sub> | 0/5* | 0/5* | 0/5* | 0/5* |
| Comp7Nstop<sub>t</sub> | 0/5* | 0/5* | 0/5* | 0/5* |

\* Significant difference compared to wild type determined by Fisher's exact test (p<0.05). ND not determined.

**Table 4.**
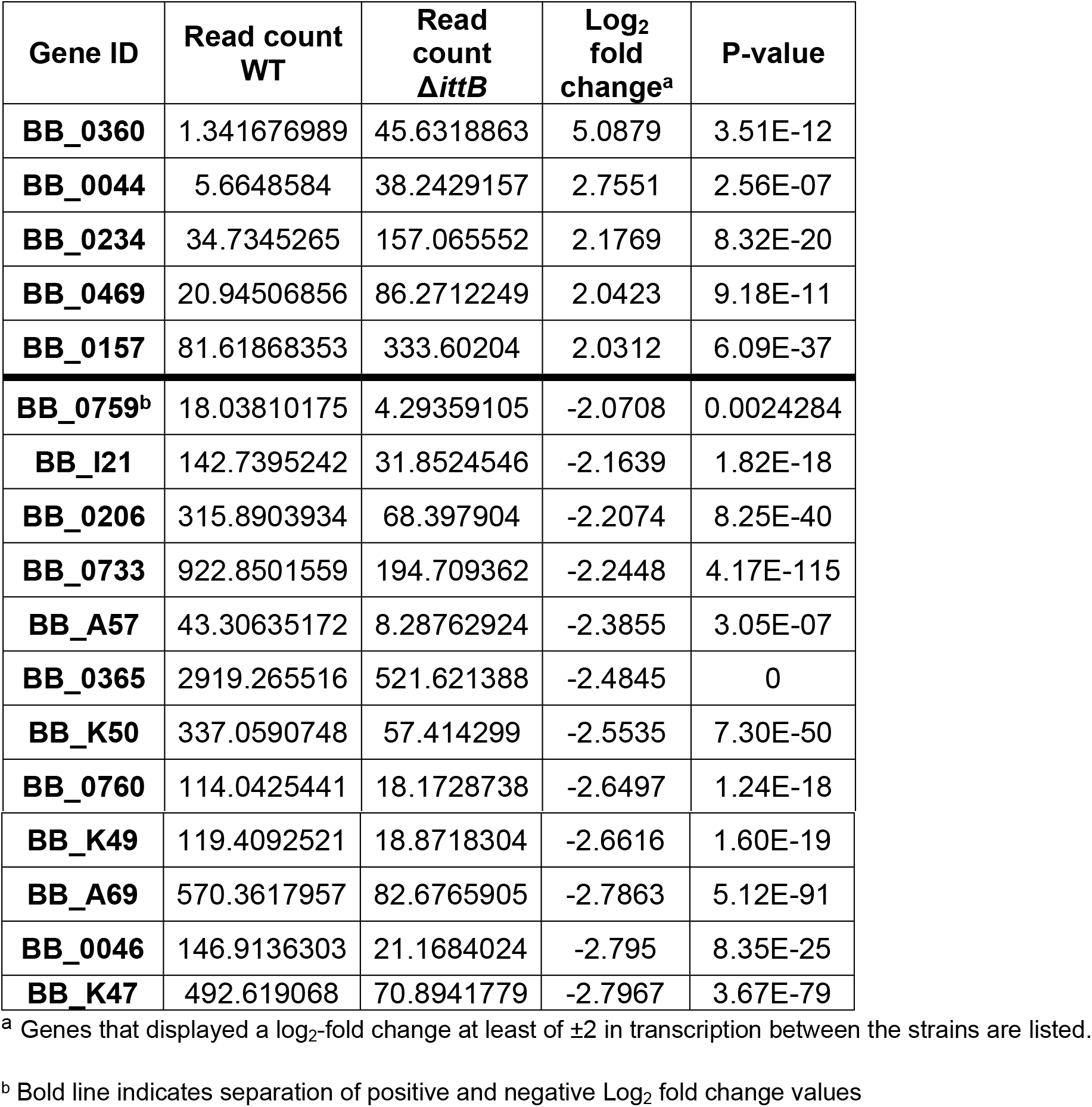
Genes differentially expressed by *ΔittB* compared to wild type *in vitro.*

## Discussion

In this study, we attempted to further characterize the *bbd07/SR0726* locus of lp17 by determining whether the functional product encoded by this intergenic region is an RNA or protein, and assessing any potential regulatory effects resulting from its targeted deletion. Our findings show that genetically complemented strains of the Δ1-7 mutant featuring mutations aimed at eliminating potential protein translation (Comp7stop_*c*_ and Comp7Δstart_*c*_) colonized heart tissue at levels comparable to the wild type and native *in cis* complement (Comp7N_*c*_). This evidence strongly suggests that the *bbd07* locus encodes an sRNA and not a protein. Because the phenotype associated with deletion of this region involves attenuated tissue colonization, we elected to denote this sRNA as *ittB* (*/*nfection associated and *t*issue *t*ropic sRNA *B*) in order to remain consistent with the naming scheme used for *ittA,* a similar lp17-encoded sRNA reported recently by other investigators (44).

We reasoned that a regulatory sRNA may have drastic effects on multiple bacterial characteristics if it were transcribed from a high-copy plasmid, and that investigation of these effects could provide insight into the native function of *ittB*. Indeed, spirochetes overexpressing *ittB* were unable to establish infection in C3H mice using an inoculum of up to 10^6^ spirochetes and were also found to be non-infectious in SCID mice. This is a uniquely severe phenotype, as many sRNAs of other of pathogenic bacteria have been observed to carry subtle deletion or overexpression phenotypes, due in part to functional redundancy and incomplete inhibition/activation of target mRNA expression (30, 33–35, 49). The severity of phenotypes associated with *ittB* deletion and overexpression suggest that this sRNA may serve a key regulatory function critical for mammalian host adaptation. That is, it may either have robust regulatory activities on multiple target mRNAs that together confer fitness advantage and tissue colonization capacity, or it may regulate a single mRNA target early in a regulatory pathway, causing multiple and critical downstream regulatory effects. In either case, it appears that the effects of the absence or overexpression of *ittB* cannot be fully overcome *in vivo* by compensatory or redundant regulatory mechanisms.

Recently, it was shown that the Δ1-7 mutant displays a dysregulated antigenic proteome during murine infection (40). Specifically, two proteins appeared to be derepressed in the absence of *ittB.* Based on this, the lack of infectivity exhibited by Comp7N_*t*_ seemed likely to be the result of antigen dysregulation during *in vitro* growth and inoculum preparation, and perhaps over-repression of protein(s) required for infection in the pre inoculum. Unexpectedly, however, *ittB* overexpression (Comp7N_*t*_) appeared to have no effect on antigen expression *in vitro.* One possible explanation for this outcome is that the lack of infectivity exhibited by Comp7N_*t*_ spirochetes is not due to altered antigen expression, but rather in the expression of cytosolic proteins such as those involved in metabolism. Alternatively, it may be that altered antigens are only detectable in spirochetes when propagating within the infected host environment. Indeed, we observed *ittB* to be transcribed roughly 100-fold higher by wild type spirochetes colonizing heart tissue compared to *in vitro*-grown spirochetes. Even higher transcription levels were detected in bladder and joint tissues. Thus, analysis of spirochetes cultivated within the host will likely be required to properly determine any gene dysregulation by the *ittB* mutant clone.

Through targeted deletion of the 317 bp intergenic region of lp17 encoding *ittB*, we were able to verify that it results in an infectious phenotype similar to the Δ1-7 mutant clone. Most importantly, Δ*ittB* spirochetes were not recovered from heart tissue of any infected mice. The altered *in vivo* antigenic profile of the Δ1-7 mutant demonstrated in a previous study was also reproduced in the current work with the newly generated Δ*ittB* mutant. Global transcriptomic analysis of the effects of *ittB* deletion revealed potential insight into the severe phenotypes associated with *ittB* deletion and overexpression (Δ1- 7 and Comp7N_*t*_, respectively). Specifically, *bb0360* was found to be approximately 34-fold more abundant in spirochetes lacking *ittB.* While this gene is annotated as a hypothetical protein with no predicted function, it has been shown to be co-transcribed with four other genes *(bb0360-bb0364).* In that study, *bb0363* was revealed as a functional cyclic-di-GMP phosphodiesterase required for optimal motility and infectivity (48). *bb0360* was the only transcript of this operon that was differentially detected in our experiment. *bb0733,* however, which was also highly differentially detected (5.5-fold less in *ΔittB),* is a c-di-GMP binding protein (50). In *B. burgdorferi,* c-di-GMP serves as a cyclic nucleotide second messenger that promotes expression of tick phase-specific genes through activation of the HK1-Rrp1 regulon, which is critical for spirochete survival in ticks (51). In turn, repression of these genes is essential for spirochete survival during transmission and mammalian infection. Considering the combination of previous observations and those reported herein, it could be that *ittB* acts to repress expression of proteins produced by spirochetes during the tick- and transmission-phases of the enzootic cycle. Further work is required to solidify a regulatory connection between *ittB* and metabolism of this important molecule.

Many knowledge gaps remain for the *ittB* sRNA. Among these is the exact sequence of the transcribed RNA product. Numerous attempts thus far to solve the 5’ transcriptional start site through RACE have failed. Additionally, we have preliminary evidence that the start and stop positions reported by Popitsch *et al.* fall within a larger region that can transcribed from reproducible RT-PCR experiments, indicating that those positions are imprecise (36). One possibility for the difficulty of solving this transcript is the predicted robust secondary structure of the sRNA that may be important for its function.

To date, lp17 has been a largely understudied component of the *B. burgdorferi* genome. However, recent findings have indicated that this plasmid may serve as a regulatory hub encoding multiple factors, including protein and sRNA molecules that interact with the regulatory cascade required for this pathogen to transit through the enzootic cycle. As techniques for studying sRNA biology in bacteria continue to emerge and evolve, it is likely that further study of lp17-encoded sRNAs will reveal new mechanisms of gene regulation that will expand our understanding of this steadily emerging pathogen. This and other recent studies are elucidating the critical nature of lp17 and sRNA activities to the fitness of the Lyme disease pathogen in each environment it encounters in nature. Further research in this area holds the promise of revealing enigmatic mechanisms of Lyme disease pathogenesis that may serve as points of opportunity for the prevention of this debilitating disease.

## Material and methods

### Bacterial strains and culture conditions

The *B. burgdorferi* isolate B31-5A4 used in this work has been characterized in previous studies ((40, 52); see Table 1). *B. burgdorferi* clones were grown at 35°C under 1.5% CO_2_ in modified Barbour-Stoenner-Kelly medium (BSK-II) supplemented with 6% rabbit serum (Cedarlane) (53). Mutant strains were grown in BSK-II supplemented with kanamycin (200 μg/ml) and/or gentamicin (100 μg/ml) as indicated. Culture density was monitored by visualization under dark-field microscopy using a Petroff-Hausser counting chamber.

### Mutant and complement strain generation and screening

A list of mutant strains and their corresponding genotypes are provided in Table 1. All transformations were performed using electrocompetent *B. burgdorferi* cells as previously described (21, 54). Transformations were recovered in drug-free media for 24 hours and plated by limiting dilution with antibiotic selection to isolate clonal transformants. Positive clones were identified by PCR for introduced sequence. *In trans* complement strains were further verified by visualization of the respective shuttle vectors on agarose gel. Endogenous plasmid content was analyzed in all strains by multiplex PCR using primers specific for regions unique to each plasmid as previously described (43).

### Ethics Statement

The experiments on mice were carried out according to the protocols and guidelines approved by American Association for Accreditation of Laboratory Animal Care (AAALAC) and by the Office of Campus Veterinarian at Washington State University (Animal Welfare Assurance A3485-01 and USDA registration number 91-R-002). These guidelines are in compliance with the U.S. Public Health Service Policy on Humane Care and Use of Laboratory Animals. The animals were housed and maintained in an AAALAC-accredited facility at Washington State University, Pullman, WA. The Washington State University Institutional Animal Care and Use Committee approved the experimental procedures carried out during the current studies.

### Infection, recovery, and quantification of *B. burgdorferi* from mice

Four-week-old male immunocompetent C3H/HeJ (C3H) and immunodeficient *C3SnSmn.CB17-Prkdc^scid^/J* (SCID) (Jackson, Bar Harbor, ME) mice were infected by subcutaneous needle inoculation (near the base of the tail, to the right of the midline) with 100 μl BSK-II containing 5×10^3^ total *B. burgdorferi* cells unless indicated otherwise.

Blood (day 7 post infection), ear (day 14, 21 and 28 post infection), joint, heart, and bladder (day 28 post infection) tissues were collected and cultured in modified BSK-II supplemented with 20 μg/ml phosphomycin, 50 μg/ml rifampicin and 2.5 μg/ml amphotericin-B. Dark-field microscopy was used to determine the presence or absence of viable spirochetes for each cultured tissue sample. A sample was deemed negative if no spirochetes could be seen in ten fields of view after three weeks of culture.

For bacterial burden quantification, heart, bladder, joint, and ear tissues were harvested at day 28 post infection and immediately snap-frozen in liquid nitrogen prior to storage at −80°C. DNA was extracted using DNeasy Blood and Tissue Kit (Qiagen, Valencia, CA) following the manufacturer’s instructions. DNA samples were then cleaned and concentrated using Genomic DNA Clean & Concentrator Kit (Zymo Research, Orange, CA). Quantitative PCR for the *B. burgdorferi flaB* and mouse *β-actin* genes was performed in duplex on each sample in triplicate using TaqMan probes and the droplet digital PCR system (ddPCR^TM^, BioRad, Carlsbad, CA). All steps, including droplet generation, thermocycling, and droplet reading were performed following the manufacturer’s instructions. All reactions were performed with 4ng of template DNA. Primers and probes used for this assay are listed in Table 1.

### qRT-PCR analysis

For RNA extraction from *in vitro* cultures, triplicate cultures of wild-type, Δ1-7, Comp7N_c_ and Comp7N_t_ were grown to late log phase (1×10^8^ spirochetes ml^-1^), pooled, and pelleted by centrifugation. RNA was extracted using a hot phenol method described previously (36, 37, 55, 56). For RNA isolation from mouse tissues, three 4-6-week-old C3H mice were infected with a total of 10^5^ wild type spirochetes. Infection was verified and monitored as described above. At day 21 post infection, heart, bladder, and joint tissue were harvested and snap frozen in liquid nitrogen. Tissues were homogenized by mortar and pestle under liquid nitrogen. Homogenates were suspended in Trizol reagent (Invitrogen), and RNA was extracted following manufacturer’s instructions using chloroform followed by alcohol precipitation.

Genomic DNA was removed from RNA samples using the Turbo DNase free kit (Invitrogen). cDNA was synthesized from 1μg of RNA using the iScript cDNA synthesis kit (BioRad). Non-quantitative control PCR reactions were performed using RNA and genomic DNA templates corresponding to each sample to verify successful removal of contaminant DNA, primer specificity, and the absence of *bbd07* amplification from Δ1-7 derived DNA. A total of 1uL of cDNA from each sample was diluted 1:10 and added to quadruplicate PCR reactions. PCR was performed using Evagreen supermix and the QX200 ddPCR system according to the manufacturer’s instructions (BioRad). Reactions for each sample were performed at least in quadruplicate.

### Western blot analysis

Cultures of indicated strains were grown in triplicate to late log-phase (1×10^8^ spirochetes mL^-1^) under standard culture conditions as described above. Replicate cultures were pooled and pelleted by centrifugation, washed, and resuspended in 90μL PBS. Lysis was performed by adding 30μL of 4X Laemmli loading dye containing 10% β- mercaptoethanol, and then heating the samples to 80°C for 10 minutes following at least one freeze-thaw cycle. Lysates (10^9^ cells) were electrophoresed on a precast 4-15% polyacrylamide gradient gel (Bio-Rad). Western blotting was performed as described previously (52), using sera harvested from C3H mice infected with 5×10^3^ spirochetes of the indicated strain for 28 days. Purified polyclonal antibodies (Rockland) were used in anti-FlaB Western blots.

### RNA-seq

Cultures were grown in triplicate under standard culture conditions to mid-log phase (5×10^7^ spirochetes mL^-1^), then temperature shifted by 1:100 dilution into room temperature BSK-II. Cultures were allowed to grow to mid log phase, at which point they were subcultured again 1:100 into warm BSK-II. These final cultures were then allowed to grow under standard conditions to late log-phase (1×10^8^ spirochetes mL^-1^). Cultures were pooled, and RNA was extracted using a hot phenol method described previously (36, 37, 55, 56). After DNase treatment, RNA was assessed for integrity using an Agilent 2100 Bioanalyzer. Clean, high quality RNA samples were shipped to the Novogene Corporation for library preparation, sequencing, and data analysis (novogene.com). Briefly, 3 μg RNA was used as input material for each sample. Sequencing libraries were generated using NEBNext^®^ UltraTM Directional RNA Library Prep Kit for Illumina^®^ (NEB, USA). PCR was performed with Phusion High-Fidelity DNA polymerase, Universal PCR primers and Index (X). Paired-end sequencing was performed on the Illumina NovaSeq 6000 platform.

### Statistical analyses

Fisher’s exact test was used to determine significant differences in the ability to recover recombinant strains by culturing of tissues compared with wild type *B. burgdorferi.* Student’s t-test was performed to determine significant differences in spirochete burden in heart tissue samples from qPCR analyses, where the average burden in heart tissues from mice infected with a given recombinant strain was compared to that from mice infected with the wild type. Student’s t-test was also used to determine significant differences in the average *in vitro ittB* transcription levels between a given recombinant strain and the wild type by qRT-PCR. One-way ANOVA followed by all pairwise multiple comparison (Holm-Sidak) was used to determine significantly different levels of average *in vivo ittB* transcription by the wild type between each tissue tested.

## Acknowledgements

We thank Danny Powell, Preeti Singh, and Jessica Wong for thoughtful and critical review of the manuscript.

## Author Contributions

MAC- Conceptualization, Formal analysis, Investigation, Methodology, Validation, Writing – original draft, Writing – review and editing.

TB- Conceptualization, Formal analysis, Funding acquisition, Methodology, Supervision, Writing- review and editing.

